# Diffusional Kurtosis Imaging in the Diffusion Imaging in Python Project

**DOI:** 10.1101/2021.03.04.433972

**Authors:** Rafael Neto Henriques, Marta Correa, Maurizio Maralle, Elizabeth Huber, John Kruper, Serge Koudoro, Jason Yeatman, Eleftherios Garyfallidis, Ariel Rokem

## Abstract

Diffusion-weighted magnetic resonance imaging (dMRI) measurements and models provide information about brain connectivity and are sensitive to the physical properties of tissue microstructure. Diffusional Kurtosis Imaging (DKI) quantifies the degree of non-Gaussian diffusion in biological tissue from dMRI. These estimates are of interest because they were shown to be more sensitive to microstructural alterations in health and diseases than measures based on the total anisotropy of diffusion which are highly confounded by tissue dispersion and fiber crossings. In this work, we implemented DKI in the Diffusion in Python (DIPY) project - a large collaborative open-source project which aims to provide well-tested, well-documented and comprehensive implementation of different dMRI techniques. We demonstrate the functionality of our methods in numerical simulations with known ground truth parameters and in openly available datasets. A particular strength of our DKI implementations is that it pursues several extensions of the model that connect it explicitly with microstructural models and the reconstruction of 3D white matter fiber bundles (tractography). For instance, our implementations include DKI-based microstructural models that allow the estimation of biophysical parameters, such as axonal water fraction. Moreover, we illustrate how DKI provides more general characterization of non-Gaussian diffusion compatible with complex white matter fiber architectures and grey matter, and we include a novel mean kurtosis index that is invariant to the confounding effects due to tissue dispersion. In summary, DKI in DIPY provides a well-tested, well-documented and comprehensive reference mplementation for DKI. It provides a platform for wider use of DKI in research on brain disorders and cognitive neuroscience research. It will ease the translation of DKI advantages into clinical applications.

## 1 INTRODUCTION

Diffusion-weighted magnetic resonance imaging (dMRI) uses a pair of directional gradient pulses to induce rephasing of proton spins, which depends on the motion of water molecules within each measurement voxel (Stejskal and Tanner, 1965; Le Bihan and Breton, 1985). Although dMRI measurements are typically made within voxels on the order of millimeters in size, they provide a view into the microstructural properties of human tissue *in vivo*. This is because the image contrast provided by dMRI is sensitive to micron-scale distances that are probed through the random motion of water within a small amount of time between the two gradient pulses (Kiselev, 2021).

The dMRI signal in each voxel is typically approximated as a three-dimensional Gaussian distribution (Basser et al., 1994; Le Bihan and Johansen-Berg, 2012), by estimating a 2^*nd*^ order tensor in every voxel. In addition to the directional information about the principal diffusion direction of the Gaussian distribution, which can be used for tractography (Mori et al., 1999; Jones, 2008), this 2^*nd*^ order tensor can be used to extract scalar measures, such as the mean diffusivity (MD) and the diffusion fractional anisotropy (FA) (Basser and Pierpaoli, 1996). The diffusion tensor imaging (DTI) model provides both an accurate fit to the dMRI signal in a wide range of experimental conditions (Rokem et al., 2015), as well as useful information to probe tissue maturation or degeneration (e.g., Pfefferbaum et al. (2000); Moseley (2002); Lebel and Beaulieu (2011); Le Bihan and Johansen-Berg (2012)). Indeed, the diffusion tensor model is so commonly used in practice that dMRI is sometimes also referred to as DTI. However, in many important cases, it is known to be systematically biased (Jones et al., 2013; De Santis et al., 2014; Henriques et al., 2015). The bias arises because, while the Gaussian model can be an adequate description for diffusion of a population of water molecules that are experiencing the same degree of restriction in a single tissue compartment type, it does not fully represent the diffusion properties of multiple different populations of water molecules inside complex biological tissue (Frank, 2001a,b). Consequently, DTI provides measures that are not only sensitive to microstructural properties but also to confounding factors such as the orientation dispersion of tissue components (Wheeler-Kingshott and Cercignani, 2009; Henriques et al., 2015), as well as the parameters of the MRI acquisition (Jones and Basser, 2004).

As an attempt to overcome these limitations of DTI, several mechanistic models directly relate diffusion properties with specific microstructural features (e.g., (Assaf and Basser, 2005; Jespersen et al., 2007; Fieremans et al., 2011; Zhang et al., 2012; Nilsson et al., 2012)), but recent studies showed that improper assumptions can compromise the validity of these models (Henriques et al., 2019; Lampinen et al., 2019, 2017). To avoid misleading interpretation, a complete characterization of water diffusion in biological tissues can be obtained using phenomenological models, which are also known as signal representation techniques (Novikov et al., 2018a). Diffusional kurtosis imaging (DKI) is a signal representation model that directly estimates the degree to which water diffusion deviates from a single Gaussian component (Jensen et al., 2005). Describing water diffusion in every voxel as an infinite mixture of Gaussian components, rather than a single Gaussian, the excess-kurtosis measured by DKI can be directly related to the variance of apparent diffusivities across different tissue components (Jensen et al., 2005; Jensen and Helpern, 2010; Fieremans et al., 2011). DKI is also sensitive to non-Gaussian diffusion effects due to the interaction of water molecules with boundaries (e.g., collisions with cell membranes or myelin sheaths) and obstacles (e.g. organelles, macromolecules) of different intra- and extra-cellular compartments macromolecules (Callaghan et al., 1991; Paulsen et al., 2015; Jespersen, 2018; Dhital et al., 2018; Henriques et al., 2020a). This means that the scalars provided by DKI closely relate to microstructural alterations in health and in brain diseases (Hui et al., 2012; Fieremans et al., 2013; Benitez et al., 2014; Marrale et al., 2016; Huber et al., 2019) and has led to the development of extensions of DKI that provide inferences about specific aspects of the microstructure (Fieremans et al., 2011; Jespersen, 2018,?; Novikov et al., 2018b; Henriques et al., 2019). In addition, DKI provides information about the diffusion orientation distribution function (dODF) within a voxel, that can be used for tractography (Lazar et al., 2008; Jensen et al., 2014; Henriques et al., 2015; Glenn et al., 2015a, 2016).

Despite the utility of DKI, the use of these analysis methods in practice depends on the availability of a free and open-source implementation of the methods. The present paper discusses the implementation of the DKI model in the DIPY project. DIPY is an open-source software library that provides implementations of many different methods for analysis of dMRI data (Garyfallidis et al., 2014). The library, implemented in the Python programming language, relies on the robust ecosystem of scientific computing tools in Python (Perez et al., 2011). It has been in continuous development since 2009, and provides a wide array of computational neuroanatomy methods. In particular, the library provides a uniform programming interface to many different dMRI signal reconstruction models and models for inferring microstructure. Here, we will focus on the modeling of the diffusion signal from individual MRI voxels using DKI, and derived microstructural models. In addition to the implementation details, we will also illustrate the advantages and drawbacks of the DKI model based on numerical simulations, and demonstrate the range of functionality implemented on openly available dMRI datasets.

## 2 METHODS

### 2.1 Theory and Implementation

Because DKI is a direct extension of the DTI model, we begin with a brief explanation of DTI and establish our notation based on this explanation. In terms of implementation, DIPY provides a class hierarchy for diffusion reconstruction models, and the DKI implementation uses components of the DTI implementation through inheritance. The code implementations are part of the DIPY source-code available at https://github.com/dipy/dipy. In addition, software implementations of the scripts and Jupyter notebooks used to generate the results and figures presented here are available at https://github.com/dipy/dipy-dki-paper.

#### 2.1.1 Diffusion Tensor Imaging

The diffusion tensor imaging (DTI) model describes the dMRI signal *S*(**n***, b*) using a 2^*nd*^ order diffusion tensor (Basser et al., 1994). In Einstein’s summation convention, the DTI model can be expressed as:

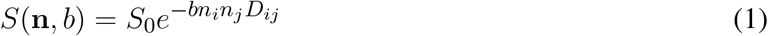

where **n** is the direction of the diffusion gradient **n** = [*n*_1_, *n*_2_, *n*_3_], *b* is a value that summarizes the intensity of diffusion weighting (calculated both from the intensity of the gradients that were applied, as well as the duration of the gradients and the interval between them), **D** is the *diffusion tensor*. This is a symmetrical matrix, which describes the variances and co-variances of the Gaussian diffusion distribution.

To solve for **D**, it is usual to normalize the direction- and b-value-specific signal by the non-diffusion-weighted signal and re-represent this ratio in the log domain:

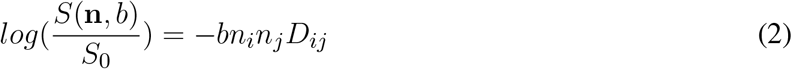

This equation can be described through a set of linear equations and solved for the six independent parameters in **D**(which requires data for at least six gradient directions (Basser et al., 1994)). In DIPY, the default fitting approach is based on a conventional weighted-least squares technique (WLS) (Chung et al., 2006). Other DTI fitting techniques are available in DIPY such as ordinary least squares (OLS), non-linear least square (NLS) and the robust estimation of tensors by outlier rejection (RESTORE) technique (Jones and Basser, 2004; Chang et al., 2005).

#### 2.1.2 Diffusion Tensor Metrics

After DTI fitting, the tensor **D** can be decomposed into three eigenvectors (**e**_1_, **e**_2_, and **e**_3_ and their respective eigenvalues (*λ*_1_ *λ*_2_ *λ*_3_) (Basser and Pierpaoli, 1996). These eigenvalues are used to compute rotationally-invariant metrics (i.e., measurements that are independent of the applied gradient direction). For instance, the mean, radial and axial diffusivities can be computed as *MD* = (*λ*_1_ + *λ*_2_ + *λ*_3_)*/*3, *RD* = (*λ*_2_ + *λ*_3_)/2, and *AD* = *λ*_1_. The eigenvalues of the diffusion tensor can also be used to produce measures of the degree of diffusion anisotropy (Basser and Pierpaoli, 1996). One of the most used diffusion anisotropy measures is the fractional anisotropy which is defined as (Basser and Pierpaoli, 1996; Glenn et al., 2015b):

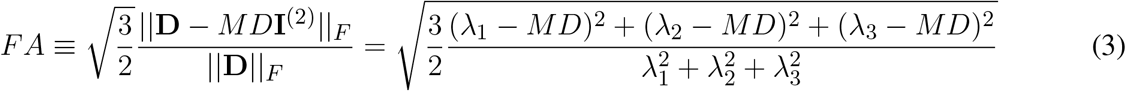

where **I**^(2)^ is the fully symmetric rank-2 isotropic tensor and ‖…‖*_F_* is the Frobenius norm of a tensor with rank *N* (Glenn et al., 2015b). The factor 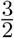 is introduced so that FA values range between 0 to 1 (from lower to higher degrees of anisotropy).

#### 2.1.3 Diffusional Kurtosis Imaging

To extend DTI and account for the excess diffusional kurtosis, DKI models the diffusional kurtosis tensor **W** in addition to the diffusion tensor **D** (Jensen et al., 2005). The DKI model can be derived by expanding the cumulants of the diffusion-weighted signal up to the 2^*nd*^ order in b (Jensen et al., 2005). Using the same notation and conventions as in Eq. 2:

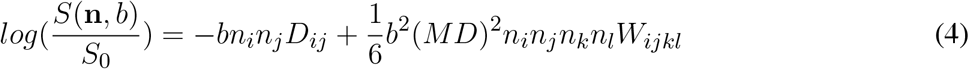

Similar to DTI, the DKI model can also be described through a set of linear equations and solved for the six independent parameters of **D** and fifteen independent parameters of **W**, noting that **W** is axially symmetric (Tabesh et al., 2011; Lu et al., 2006). In addition to 15 different gradient directions to resolve the anisotropic information of **W**, the DKI model requires at least three b-values (these can include signals for *b*=0 in addition to two non-zero b-values). In DIPY, the default DKI fitting was implemented based on a weighted-least squares (WLS) technique in which weights are defined from previous diffusion parameter estimates (Veraart et al., 2013). This fitting approach was shown to provide diffusion and kurtosis estimates with higher accuracy and precision when compared to other linear least square fitting strategies and provides faster fits when compared to non-linear least square approaches. Additionally, the fitting approaches implemented in DTI (OLS, NLS, RESTORE) were adapted to the DKI model and can also be used in DIPY as alternative DKI fitting strategies.

#### 2.1.4 Kurtosis Tensor Metrics

Since it also fits the diffusion tensor, DKI can be used to estimate all DTI metrics (e.g. MD, RD, AD, FA explained above). Additionally, rotationally-invariant measures can be defined from the kurtosis tensor. In analogy to the definition of MD, the **mean kurtosis** (MK) is defined as the average of directional kurtosis coefficients across all spatial directions, which can be formulated by the following surface integral (Jensen and Helpern, 2010):

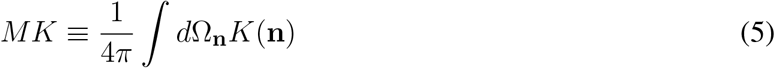

where *K*(**n**) is the directional kurtosis for a given direction **n**, which can be sampled from the fitted diffusion and kurtosis tensors as:

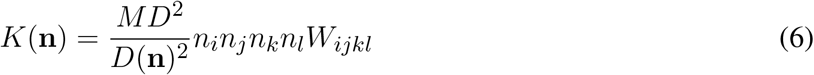

with

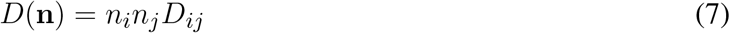

In DIPY, two approaches were implemented to compute MK:

1. the integral of equation 5 is numerically resolved by averaging directional kurtosis values sampled for a finite number of directions. Biases of discrete direction samples can be avoided by using a spherical t-design as shown by (Hardin and Sloane, 1996). For the DIPY implementation of MK, a t-design of 45 directions is used.
2. The second approach is based on the analytical solution of equation 5 (Tabesh et al., 2011), avoiding the use of discrete directional samples. This approach requires the following computing steps: a) the rotation of the DKI tensors to a frame of reference in which **D** eigenvectors are aligned to the Cartesian axis *x*, *y* and *z*. This rotated kurtosis tensor is denoted as 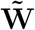; b) the evaluation of Carlson’s elliptic integrals (Carlson, 1995); and c) the treatment of the solution’s singularities for *λ*_1_ = *λ*_2_, *λ*_1_ = *λ*_3_, *λ*_2_ = *λ*_3_, and *λ*_1_ = *λ*_2_ = *λ*_3_. These steps were vectorized for optimal processing speed.

A comparison of these two approaches is presented in supplementary notebooks available at https://github.com/dipy/dipy-dki-paper.

Since the directional kurtosis coefficient for a given direction **n** depends on both diffusion and kurtosis tensors (equation 6), MK as defined by Eq. 5 depends on both diffusion and kurtosis tensors. To have a mean kurtosis metric independent to the diffusion tensor, the **mean kurtosis tensor** (MKT) is defined as (Hansen et al., 2013; Hansen and Jespersen, 2017):

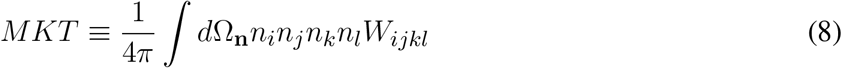

This latter quantity can be directly computed from the trace of the kurtosis tensor:

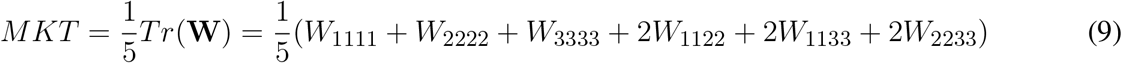

Although (Hansen et al., 2013) showed that MKT and MK have similar contrasts in neural tissue, the mean kurtosis tensor was also implemented in DIPY by applying equation 9 to **W** directly.

For voxels containing well-aligned structures, the **radial kurtosis** is defined as the average of the directional kurtosis across all directions perpendicular to the main direction of fibers which should correspond to the diffusion tensor main direction **e**_1_) (Jensen and Helpern, 2010; Tabesh et al., 2011):

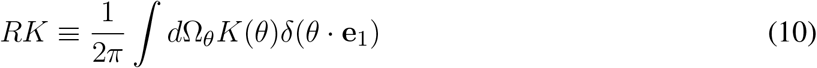

Similar to the estimation of MK, DIPY provides two methods to compute RK based on a numerical and an analytical approach:

1. Eq. 10 can be numerically computed by averaging directional kurtosis values for directions perpendicular to **e**_1_. The directional kurtosis values can be sampled from the fitted kurtosis tensor using equation 6. Directions *θ* that are evenly sampled and perpendicular to **e**_1_^1^.
2. Alternatively, Eq.4 can be solved analytically, avoiding discrete perpendicular direction samples (Tabesh et al., 2011). This approach requires the rotation of the kurtosis tensor and the treatment of a singularity for *λ*_2_ = *λ*_3_.

The **axial kurtosis** is defined as the directional kurtosis along the main direction of well-aligned structures:

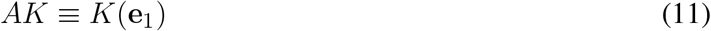

This quantity can be computed from one of the following:

1. The directional kurtosis coefficient along the tensor eigenvector (i.e. applying **e**_1_ into Eq. 6;
2. From the tensor element 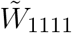 of the rotated kurtosis tensor (i.e. 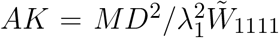, (Tabesh et al., 2011)).

Although both approaches lead to the exact calculation of AK, the former and latter estimators will be referred to as the numerical and analytical methods respectively, to keep the nomenclature consistent to the estimation strategies of MK and RK.

Similar to the definition of FA for the diffusion tensor, **the anisotropy of the kurtosis tensor** can be quantified as (Glenn et al., 2015b):

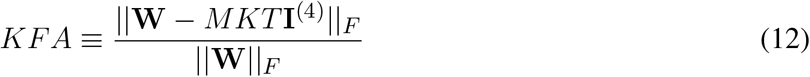

where **I**^(4)^ is the fully symmetric rank 4 isotropic tensor, ‖…‖*_F_* is the Frobenius norm (Glenn et al., 2015b), and MKT is the mean kurtosis tensor defined by Eq. 9. Analogous to the FA of the diffusion tensor, KFA quantifies lower to higher kurtosis tensor anisotropy in a range between 0 and 1.

#### 2.1.5 White Matter Tract Integrity Model

One way to interpret the information captured by DKI is to fit additional microstructural models to the diffusion and kurtosis tensors (Jensen et al., 2005; Jensen and Helpern, 2010; Fieremans et al., 2011; Novikov et al., 2018b; Jespersen, 2018). This approach provides DKI-derived scalar quantities that are potentially more specific to microstructural properties of the tissue, such as the fraction of signal contributions due to extra- or inter-cellular spaces. However, as in the case of microstructural models applied directly to dMRI signals (Jespersen et al., 2007; Zhang et al., 2012; Assaf and Basser, 2005; Kaden et al., 2016b), the interpretation of these quantities is only valid if the assumptions of the microstructural models are met (Henriques et al., 2019; Lampinen et al., 2019, 2017; Novikov et al., 2018a). The White Matter Tract Integity (WMTI) model (Fieremans et al., 2011, 2013) relates the diffusion and kurtosis tensors to the parameters of a two compartments model representing the intra- and extra-cellular components of aligned white matter fibres:

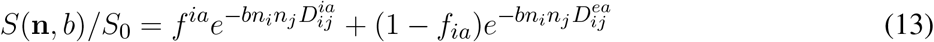

where *f^ia^* is the intra-axonal water fraction (AWF), **D**^*ia*^ is the intra-axonal diffusion tensor and **D**^*ea*^ is the extra-axonal diffusion tensor.

The WMTI model relies on the following assumptions:

1. The tissue is only described by non-exchanging intra- and extra-cellular compartments. Other signal components, such as from glia cell, have to be in fast exchange with the extra-cellular compartment.
2. The intra-cellular diameter of axons is much smaller than the volume probed by diffusing particles. That is, intra-cellular RD is practically zero.
3. In all directions water-molecules can more freely move in the extra-cellular volume. That is, intra-cellular AD is smaller than extra-cellular AD
4. Intra-cellular spaces are well-aligned to each other. This does not apply to voxels containing fiber dispersion or crossing.
5. Effects of the interaction of water molecules with the boundaries of different intra- and extra-cellular compartments (e.g., collision with cell membranes or myelin sheaths) or with macromolecules are negligible.

Despite several studies that have demonstrated that these assumptions are unlikely to hold in many cases (e.g., (Dhital et al., 2018)), WMTI measures were still shown useful as sensitive biomarkers for the characterization of progression of white matter microstructural alterations in health and disease (e.g., Hui et al. (2012); Fieremans et al. (2013).

In DIPY, WMTI is implemented as follows:

1. <u>Computing the maximum directional kurtosis</u>. Kurtosis is evaluated using Eq.6 for 100 uniformly distributed directions **n** and direction of the maximal value is used to seed a quasi-Newton method algorithm to optimize the following problem:

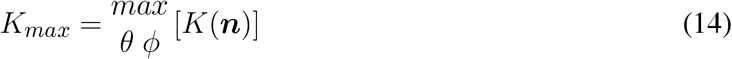

where *θ* and *φ* are the polar and azimuth coordinates of the unit direction ****n**** that maximizes the kurtosis.
2. <u>Computing the axonal water fraction</u>. For a system described by equation 5, the maximum kurtosis is expected to be perpendicular to the main direction (Fieremans et al., 2011; Henriques et al., 2015). Under the assumption that intra-cellular RD is zero, the axonal volume fraction (AWF) is computed as (Fieremans et al., 2011; Jespersen, 2018; Jensen and Helpern, 2010):

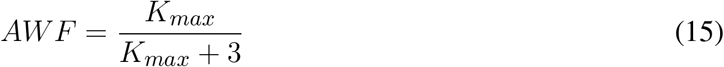
3. <u>Decoupling the compartmental diffusivities</u>. Assuming that extra-axonal diffusivity is always higher than the intra-axonal diffusivity, the directional diffusivities for both intra- and extra-cellular compartments *D*(**n**)_*i*_ and *D*(**n**)_*e*_ are estimated for given directions **n**, using the following expressions:

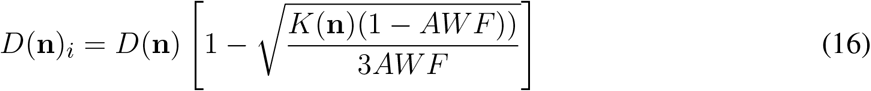

and

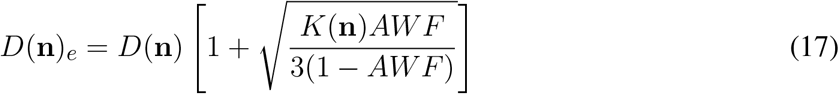

where *D*(**n**) and *K*(**n**) are computed from equations 7 and 6. The tensors **D**^*ia*^ and **D**^*ea*^ are computed from *D*(**n**)_*i*_ and *D*(**n**)_*e*_ samples for at least six different directions **n** (Fieremans et al., 2013).
4. <u>Deriving WMTI metrics</u>. In addition to the AWF, other WMTI metrics are defined from tensors **D**^*ia*^ and **D**^*ea*^: the axonal diffusivity *D^ia^* defined as the trace of **D**^*ia*^; the axial and radial diffusivities of the extra-cellular diffusion tensor *AD^ea^* and *RD^ea^*; and the extra-cellular tortuosity which is defined as the ratio between *AD^ea^* and *RD^ea^*.

After WMTI model fitting, a class instance is created containing all WMTI model parameters and derived metrics. AWF, *D^ia^*, *AD^ea^*, *RD^ea^*, and the extra-cellular tortuosity are as the respective class methods/attributes: “awf”, “axonal diffusivity”, “hindered ad”, “hindered rd”, and “tortuosity”.

#### 2.1.6 Mean Signal Diffusional Kurtosis Imaging

The DKI model aims to characterize the full 3D directional dependence of diffusional kurtosis, which is influenced by tissue microstructural properties. For example, by the sizes of different compartments, their apparent diffusivities, and volume fractions. However, directional kurtosis is also affected by tissue organization: the degree of dispersion, crossing or fanning (Henriques et al., 2015). Increased specificity toward microstructural properties can be achieved by measuring a scalar excess-kurtosis index from powder-averaged signals (Henriques, 2018; Henriques et al., 2019). That is, averaged signals across evenly-sampled gradient direction for each b-value that are independent of the tissue orientation distribution function (Kaden et al., 2016b; Jespersen et al., 2013). In DIPY, this technique is referred to as the mean signal diffusional kurtosis imaging (MSDKI). Analogous to the derivation of DKI from the directional diffusion-weighted signals, the MSDKI model can be derived from powder-averaged signals 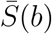 using the second-order cumulant expansion (Henriques, 2018; Henriques et al., 2019):

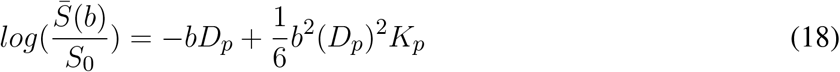

where *D_p_* and *K_p_* are the diffusivity and excess-kurtosis of the powder-average signals. In DIPY, these quantities are referred to as the mean signal diffusivity (MSD) and mean signal kurtosis (MSK). It is important to note that, while MSD theoretically is equal to the standard mean diffusivity (MD) (Henriques et al., 2019), MSK is equal to mean kurtosis tensor (MKT) subtracted by a mesoscopic dispersion correction factor ψ; which can be calculated from the diffusion tensor (Henriques et al., 2020a), i.e:

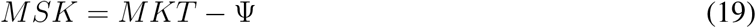

with

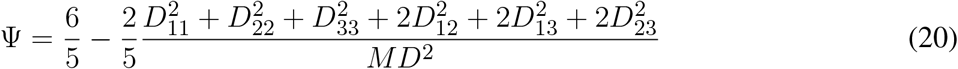

Diffusion-weighted data can be acquired with different numbers of gradient directions *N_g_* for different b-values. Therefore, in DIPY, the MSDKI model (eq. 18) is fitted using a weighted least square approach in which weights for each b-value are set to 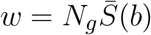 (Henriques, 2018).

#### 2.1.7 MSDKI-based Microstructural models

Analogous to the DKI metrics, the parameters of MSDKI can also be related to microstructural models. For instance, Henriques et al. (2019) showed that MSDKI captures the same information than the spherical mean technique (SMT) microstructural models (Kaden et al., 2016b,a). In this way, the SMT model parameters can be directly computed from MSDKI. In DIPY, the intrinsic diffusivity (*D_I_*) and the axonal water fractions (AWF) of the two-compartmental SMT model parameters (Kaden et al., 2016a) can be estimated from MSDKI parameters was implemented by inverting the following equations (Henriques et al., 2019):

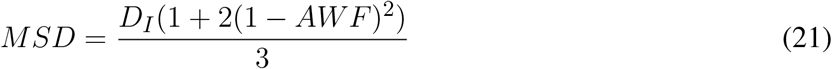

and

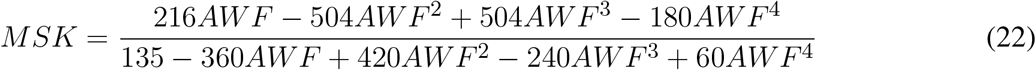

Although SMT models are general to any tissue configuration (i.e. general to well-aligned, crossing or dispersing fibers), the two-compartmental model assumes that: 1) the axial diffusivities of both intra- and extra-cellular spaces are equal to the in the intrinsic diffusivity (*D_I_*), 2) the extra-cellular radial diffusivity follows the first order tortuosity assumption (*RD* = (1 *AWF*)*D_I_*); and 3) the intra-cellular radial diffusivity is zero.

#### 2.1.8 Numerical Simulations for DKI Unit Testing

DIPY uses both rigorous unit testing (with pytest) and continuous integration (Travis, Appveyor and Azure Pipelines) to validate software implementations against known analytically-derived cases, and to assess any change to the software with fixes and enhancements that are introduced.

To test our DKI implementations, numerical simulations were produced for a sum of *N* Gaussian diffusion compartments (i.e., effects of interaction between diffusing water molecules and compartment’s obstacles are assumed to be negligible):

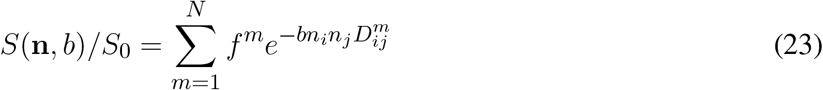

where *f^m^* is the apparent water fraction of a tissue compartment *m*, and 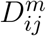 are the elements of its Gaussian diffusion. The ground truth of the elements of the total diffusion tensor *D_ij_* and total kurtosis tensor *W_ijkl_* of the multiple compartment system are computed as (Jensen and Helpern, 2010; Lazar et al., 2008; Henriques et al., 2015):

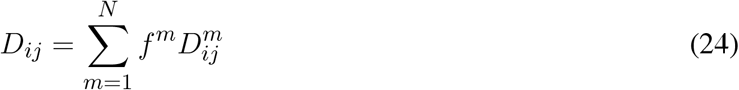

and

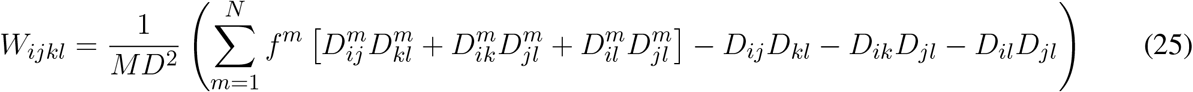

To remove the effects of the cumulants of equation 23 associated with terms higher than the 2^*nd*^ order in b, synthetic diffusion-weighted signals for implementation testing are produced by plugging the ground truth **D** and **W** tensors into equation 4. These synthetic signals were produced for different ground truth compartmental tensors **D**^(*m*)^ (with different axial and radial diffusivities **AD**^(*m*)^ and **RD**^(*m*)^ and different orientations), so that different diffusion metrics are validated for different simulation scenarios. A summary of different sets of ground truth parameters and the different checks used for DKI unit testing are presented in table 1. In addition to these, the Carlson integrals were also evaluated according to the numerical checks suggested in the original work by Carlson (Carlson, 1995).

**Table 1.** Details of numerical Simulations used for DKI Unit Testing Numerical - a brief description of each simulations, the simulation compartmental diffusion parameters (where *AD*^(*m*)^, *RD*^(*m*)^, *f*^(*m*)^, *φ*^(*m*)^ and *θ*^(*m*)^ are the axial diffusivity, radial diffusivity, water fraction, polar orientation angle and azimuthal orientation angle of a given compartment *m*), and the checks preformed are presented on the left, middle and right columns respectively.

| Simulations | G.T. Parameters | Tests |
| --- | --- | --- |
| Isotropic tensors<br>(2 compartments) | $AD^{(1)} = RD^{(1)} = 0.99$<br>$f^{(1)} = 0.5$<br>$AD^{(2)} = RD^{(2)} = 2.26$<br>$f^{(2)} = 0.5$ | 1) Check if MK, RK, AK, MKT, MSK, and directional kurtosis samples are equal to the G.T. isotropic kurtosis value of 0.4581;<br>2) Check that KFA = 0;<br>3) Check that the maximum kurtosis calculation procedure does not generate an error (note that isotropic kurtosis does not have a maximum); |
| Single fiber<br>(2 compartments)<br>(Henriques et al., 2015) | $AD^{(1)} = 0.99,$<br>$RD^{(1)} = 0,$<br>$f^{(1)} = 0.49$<br>$AD^{(2)} = 2.26,$<br>$RD^{(2)} = 0.87,$<br>$f^{(2)} = 0.51$<br>$\theta^1 = \theta^2 = rand$<br>$\phi^1 = \phi^2 = rand$ | 1) Check if rotated Kurtosis tensor $\tilde{\mathbf{W}}$ have only the following non-zero elements:<br>$\tilde{W}_{1111} = 1.7067, \tilde{W}_{2222} = \tilde{W}_{3333} = 0.800,$<br>$\tilde{W}_{1122} = \tilde{W}_{1133} = 0.3897, \tilde{W}_{2233} = 0.2670$<br>2) Test if the direction of maximum kurtosis is perpendicular to the main fiber direction (Henriques et al., 2015);<br>3) Test if the value of maximum apparent kurtosis is equal to RK;<br>4) Test if WMTI fitted parameters matches their G.T. values |
| Crossing fibers<br>(4 compartments =<br>2 intra-cellular +<br>2 extra-cellular) | $AD^{(1)} = AD^{(3)} = 0.99$<br>$RD^{(1)} = RD^{(3)} = 0$<br>$f^{(1)} = f^{(3)} = 0.245$<br>$AD^{(2)} = AD^{(4)} = 2.23$<br>$RD^{(2)} = RD^{(4)} = 0.87$<br>$f^{(2)} = f^{(4)} = 0.255$<br>$\theta^1 = \theta^2 = 80^\circ$<br>$\phi^1 = \phi^2 = 10^\circ$<br>$\theta^3 = \theta^4 = 20^\circ$<br>$\phi^3 = \phi^4 = 30^\circ$ | 1) Check if estimated $D_{ij}$ and $W_{ijkl}$ elements for all DKI fitting procedures (OLS, WLS, NLS, RESTORE) match the ground truth parameters given by equations 23 and 24;<br>2) Check if rotated Kurtosis tensor $\tilde{\mathbf{W}}$ have the same values of a reference $\mathbf{W}$ generated for a scenario in which the D is already aligned to the cartesian axis $x, y$, and $z$;<br>3) Check if MK, AK, and RK numerical solutions match their analytical solutions;<br>4) Test if the direction of maximum apparent kurtosis is perpendicular to both fibers (Henriques et al., 2015) |
| Fibers crossing at $90^\circ$<br>(4 compartments=<br>2 intra-cellular +<br>2 extra-cellular) | (same diffusivities as above)<br>$\theta^1 = \theta^2 = 90^\circ$<br>$\phi^1 = \phi^2 = 0^\circ$<br>$\theta^3 = \theta^4 = 20^\circ$<br>$\phi^3 = \phi^4 = 30^\circ$ | 1) Check if MK matches the values of the previous simulation (Note that fibers crossing at $90^\circ$ has a prolate diffusion tensor which corresponds to MK analytical solution singularity) |
| Two crossing compartments | $AD^{(1)} = AD^{(2)} = 1.7$<br>$RD^{(1)} = RD^{(2)} = 0.3$<br>$f^{(1)} = f^{(2)} = 0.5$ | 1) Check if KFA for this scenario is equal to $\sqrt{13/5}$ (Gleen et al., 2015) |

### 2.2 Simulated experiments

To illustrate the sensitivity and specificity of its different metrics, DKI is first processed on single voxel synthetic signals which were produced using equation 23 for four different sets of ground truth parameters (Figure 1A):

1. Single axial symmetric diffusion tensor component with axial and radial diffusivities of 1.7e-3 and 0.3e-3 *mm*^2^/*s*. These diffusivities were set according to typical white matter diffusion tensor estimates.
2. Two aligned axial symmetric diffusion tensor components with equal volume fractions. This scenario was produced to consider typical diffusion heterogeneity of voxels containing single healthy white matter fiber populations. As a toy-model two diffusion tensors are referred to as intra- and extra-cellular components. Axial diffusivity, radial diffusivity and volume fraction for the intra-cellular components were set to 1.4e-3 *mm*^2^/*s*, 0.1e-3 *mm*^2^/*s*, and 0.5, while the axial diffusivity, radial diffusivity and volume fraction for the extra-cellular component were set to 2e-3 *mm*^2^/*s*, 0.5e-3 *mm*^2^/*s*, and 0.5, respectively.
3. Two aligned axial symmetric diffusion tensor components with different volume fractions. This scenario was produced as a toy-model of a damaged single fibre population. For this, relative to scenario 2, the volume fraction of the intra-cellular cellular component was decreased to 0.3 while the radial diffusivity of the extra-cellular space was increased to 0.7e-3 *mm*^2^/*s*.
4. Four axial symmetric diffusion tensor components with equal volume fractions. This scenario was produced as a toy model to represent the intra- and extra-cellular contributions of two fiber populations crossing at 60 degrees.

**Figure 1.**
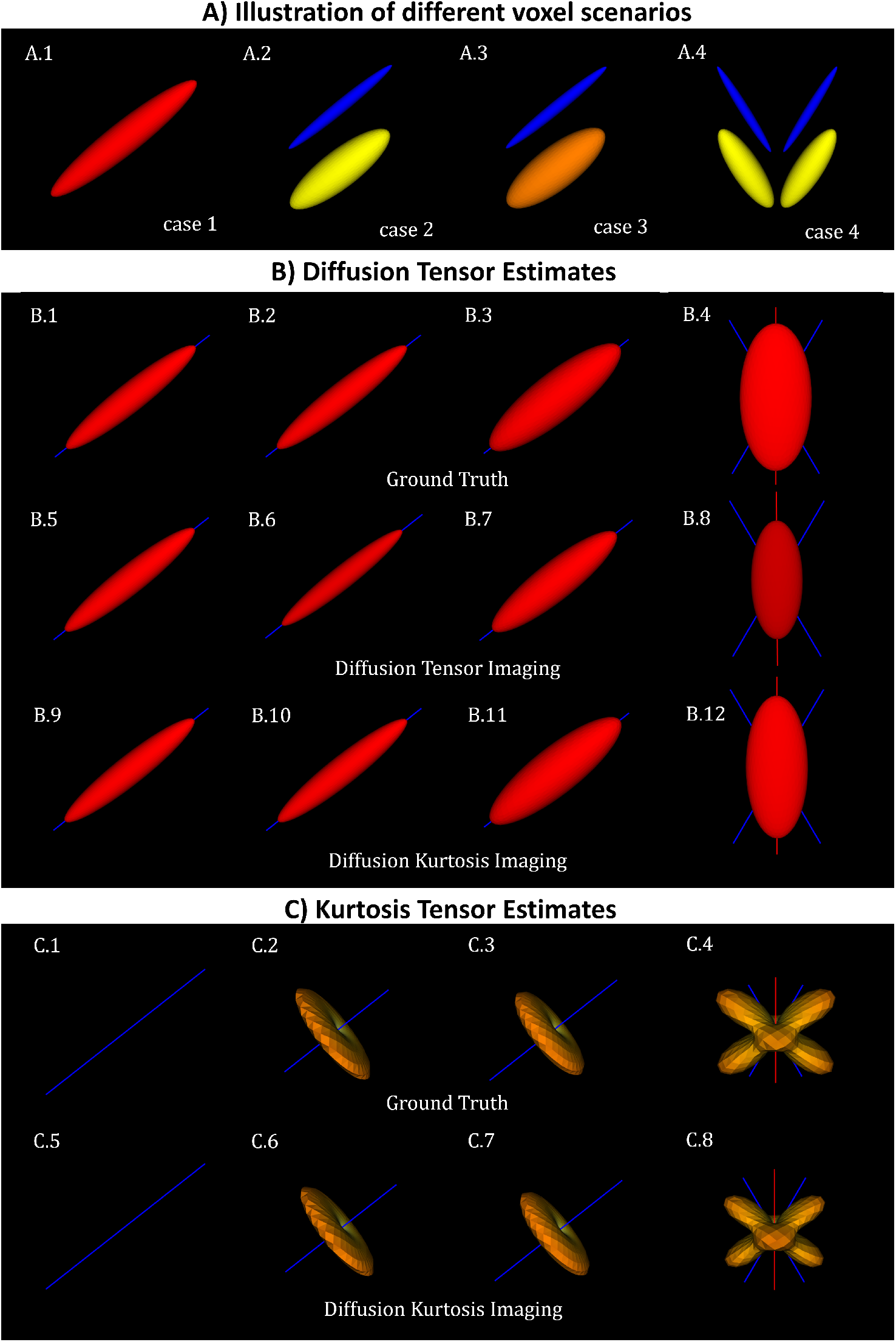
Diffusion and kurtosis tensors from single and multi-tensor toy-models. A) Illustration of the tensor components of each simulation case: (A.1) single tensor with *AD* and *RD* = 1.7×10^(^*−*3)*mm*^2^*/s* and 1.3 × 10^(^ − 3)*mm*^2^*/s*; (A.2) mixture of collinear tensors with *AD*_1_, *RD*_1_, *AD*_2_, and *RD*_2_ = 1.4 × 10^(^ − 3), 0.1 × 10^(^ − 3), 2 × 10^(^ − 3), 0.5 × 10^(^ − 3)*mm*^2^*/s* (toy-model of a healthy fiber population); (A.3) mixture of collinear tensors with *AD*_1_, *RD*_1_, *AD*_2_, and *RD*_2_ = 1 10^(^ 3), 0.1 10^(^ 3), 2 10^(^ 3), 0.7 10^(^ 3)*mm*^2^*/s* (toy-model of a damaged fiber population); and (A.4) mixture of crossing tensors (toy-model of crossing healthy fiber populations). B) Diffusion tensors for each voxel simulation: (B.1-B.4) Ground truth diffusion tensors; (B.5-B.8) diffusion tensors computed from DTI fit; and (B.9-B.12) diffusion tensors computed from DKI. C) kurtosis tensors for each voxel simulations: (C.1-C.4) Ground truth kurtosis tensors; (C.5-C.8) Kurtosis tensors fitted by DKI. In this figure, diffusion tensor are plotted in their ellipsoid representation, while kurtosis tensors are plotted as the 3D spatial variation of apparent kurtosis coefficients.

The synthetic signals for each set of ground truth parameters are generated according to the same gradient directions **n** and b-values *b* of the CFIN dataset (vide infra).

### 2.3 MRI experiments

Open-source software tools such as DIPY serve a particularly important role in advancing science in a period in which we are seeing an increase in availability of open datasets. In the work presented here, we use some of these open datasets to illustrate typical contrasts of different DKI metrics and to show the functionality of our DKI implementations.

#### 2.3.1 DKI-specific dataset

To illustrate some typical DKI contrasts, we processed a dataset that was specifically collected to support the development of DKI modeling approaches (Hansen and Jespersen, 2016a), and which can be automatically downloaded using DIPY. This data will be referred to as the **CFIN dataset**. This dataset was acquired on Siemens 3T (32 channel head coil) along 33 diffusion gradient directions for multiple b-values sampled in steps of 200 *s*^2^ from 200 *s*^2^ to 3000 *s*^2^ in addition to a single acquisition for b-value=0. All data was acquired with inversion recovery to suppress cerebrospinal fluid signal. More imaging parameters details of the CFIN dataset were previously reported (Hansen and Jespersen, 2016a).

Since DKI involves the estimation of a large number of parameters and since the non-Gaussian components of the diffusion signal are more sensitive to artefacts (Tax et al., 2015; Jensen et al., 2005), it might be favorable to suppress the effects of noise and artefacts before diffusional kurtosis fitting. Noise of the CFIN dataset was suppressed using the Marcenko-Pastur PCA denoising algorithm as proposed by (Veraart et al., 2016b), while the impacts of Gibbs ringing artefacts were attenuated using a sub-voxel Fourier Transform shifts (Kellner et al., 2016; Henriques, 2018). Both pre-processing procedures were shown to provide optimal performances for DKI (Ades-Aron et al., 2018a; Henriques, 2018). Open-source implementations of these procedures are available in DIPY.

#### 2.3.2 Testing DKI in a large open dataset

We test the DKI implementation in data from the Human Connectome Project (HCP) (Glasser et al., 2016). The HCP has collected data about brain connectivity from 1,200 individuals and includes measurements of functional and structural MRI, as well as dMRI, in addition to many measurements of phenotypical information (e.g., behavioral assessments) (Sotiropoulos et al., 2013; Glasser et al., 2016). We used data from the 2015 900-subject release to compare the DTI and DKI models. We used data from the 788 subjects for which a complete dMRI measurement was available. Briefly: the measurements conducted included 270 measurement directions, 90 directions in each of three b-value tiers: *b* ≈ 1000*s/mm*^2^, *b* ≈ 2000*s/mm*^2^ and *b* ≈ 3000*s/mm*^2^. In addition, 18 measurements with b-values close to 0 (*b* ≈ 5*s/mm*^2^ were taken. Voxel dimensions were 1.25 1.25 1.25 *mm*^3^. We used data that was preprocessed using the HCP preprocessing pipeline. Additional details of measurement and processing were previously published (Sotiropoulos et al., 2013).

The data was accessed in the Amazon Simple Storage Web Service (S3) through the AWS Open Data program. To assess the quality of fits to the data, we performed a k-fold cross-validation procedure (Rokem et al., 2015): each subject’s diffusion-weighted data was split into k parts, based on gradient directions. In each iteration of cross-validation, 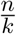 directions were left out and a model (either DKI or DTI) was fit to the remaining 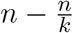 directions. The model computed was then used to predict the signal in the left out data. After k iterations of this procedure, each one of the diffusion-weighted measurements will have been predicted based on other measurements. This approach allows us to compare models with different numbers of parameters, while avoiding over-fitting to the noise in models with larger numbers of parameters, or more analytical flexibility (Stone, 1977, 1978).

Model accuracy is assessed as the coefficient of determination *R*^2^ between the predicted signals and the measured signals across the 270 diffusion-weighted measurements in each voxel in the white matter. To compare across subjects, the median *R*^2^ is computed within all of the white matter voxels in each individual. In addition to model accuracy, we can assess the precision of metrics derived from both models in this data. This is done by sub-sampling the data into different combinations of b-value tiers, as previously done in (Veraart et al., 2011). FA was calculated using DTI for the *b* ≈ 1000*s/mm*^2^ tier and for a combination of the *b* ≈ 1000*s/mm*^2^ tier and the *b* ≈ 2000*s/mm*^2^ tier. FA was also calculated using DKI for a combination of *b* ≈ 1000*s/mm*^2^ and *b* ≈ 2000*s/mm*^2^ and for a combination of *b* ≈ 2000*s/mm*^2^ and *b* ≈ 3000*s/mm*^2^. Precision is assessed by computing the difference between the two estimates of FA in each voxel in the white matter. Bias is assessed as systematic differences between the two sub-samples, while variability is assessed as the range of 95% of the differences. Small values in both of these values indicate high precision of the model estimates.

## 3 RESULTS

### 3.1 DKI simulations

The ground truth diffusion tensors of the four different voxel simulations is shown in the panel A of Fig. 1. DKI provides information about both diffusion and kurtosis tensors. For all voxel simulations, panel B of Fig. 1 shows the ground truth diffusion tensor computed using Eq. 24 (panels B.1-B.4), the diffusion tensors computed from DIPY’s DTI fit (panels B.5-B.8), and the diffusion tensors computed from DIPY’s DKI fit (panels B.9-B.12). Note that diffusion tensors for both DTI and DKI match their ground truth for voxels containing a single diffusion component (B.1, B.5, and B.9). For multi-component simulations, the diffusion tensors from DTI show, however, lower magnitudes than the ground truth tensors (B.6-B.8). On the other hand, the diffusion tensors from the DKI are closer to their ground truths (B.9-B.12). For a visualization of the kurtosis tensors, the apparent directional kurtosis computed from both ground truth (Eq. 25) and DKI-fitted tensors are shown in panel C of Fig. 1. Both ground truth and fitted kurtosis tensors are null for single diffusion component simulations (C.1 and C.5). They show maximum values perpendicular to the direction of the aligned multi-tensors (C.2-3, and C.6-7). Moreover, in contrast to the diffusion tensor, the kurtosis tensor shows to provide information on the direction of crossing multi-tensor simulations (C.4 and C.8).

Fig. 2 shows the values of diffusion and kurtosis tensor metrics obtained from DKI for all four voxel simulations. Simulation cases 1 and 2 were designed to have equal values of MD, RD, AD, and FA. However, the presence of multi-tensors for voxel case 2 is revealed by the positive values of the standard kurtosis values (i.e., MK, RK, AK). MSK is positive even for anisotropic Gaussian diffusion (case 1) because, as being a metric from directional-averaged signals, it is sensitive to diffusion variance across different gradient directions. Relative to a toy model of a healthy fiber population (voxel case 2), all standard diffusivity metrics (MD, RD and AD) show higher values for the toy model of a damaged fiber population (case 3), while all kurtosis metrics (MK, RK, AK, and MSK) shows lower values. FA shows also decreased values for the damaged fiber toy model; however, its values are even lower for the toy model of healthy crossing fibers (case 4). MSK shows equal values for simulation case 2 and 4, confirming that MSK is only dependent on the diffusion variance across voxel components and invariant to added crossing compartments.

**Figure 2.**
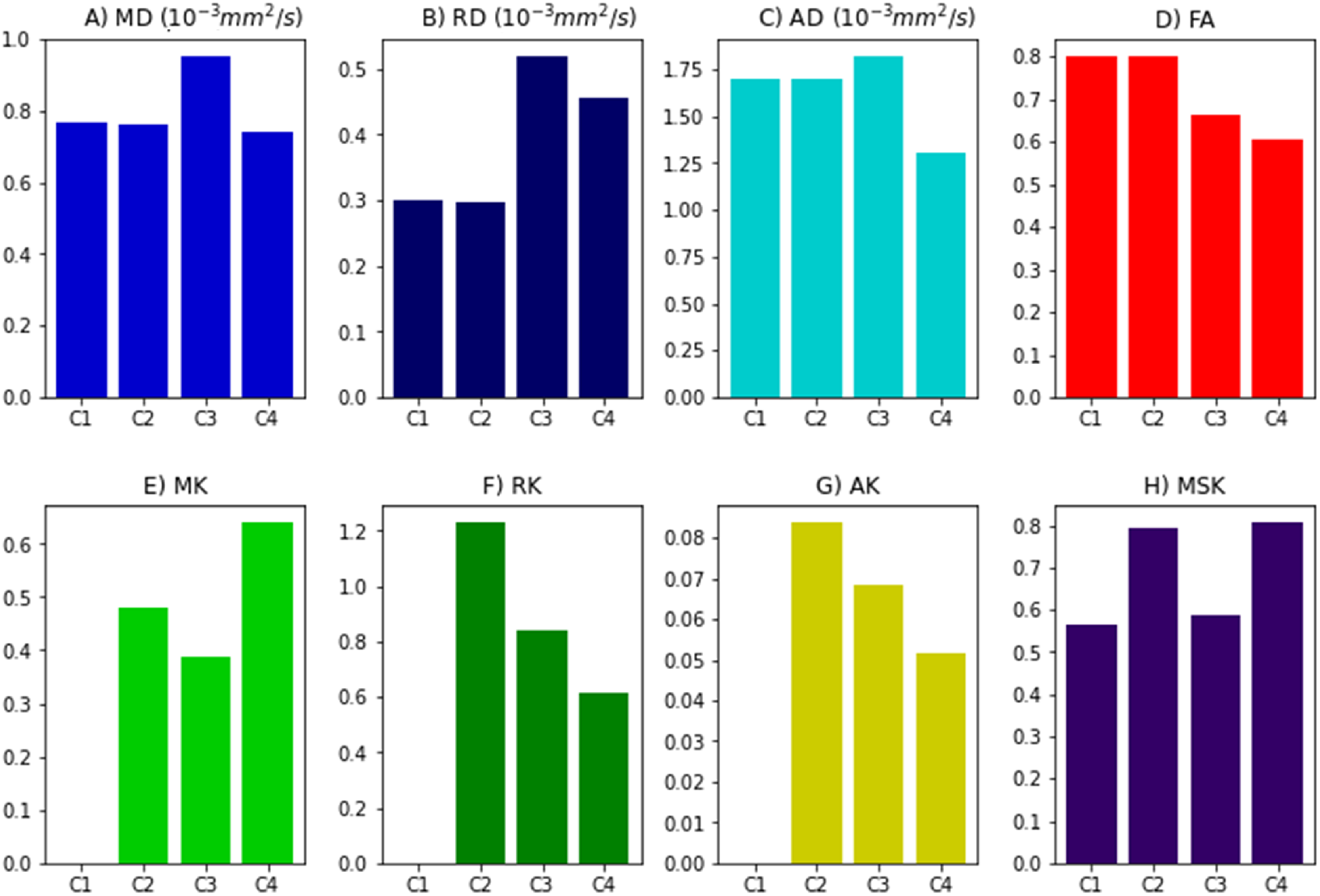
DKI and MSDKI diffusion and kurtosis metrics for the four voxel simulations: A) Mean diffusivity; B) Radial diffusivity; C) Axial diffusivity; D) Fractional Anisotropy; E) mean kurtosis; F) Radial Kurtosis; G) Axial kurtosis; H) Mean signal kurtosis.

### 3.2 Example DKI contrasts

Diffusion tensor metrics extracted from an axial slice of CFIN dataset are shown in Figure 3. Upper panels show the diffusion metrics extracted from the DTI model (Fig.3, panel A), while lower panels show the diffusion metrics extracted from the DKI model (Fig.3, panel B). MD maps show low contrast between grey and white matter (Fig.3, panel A.1 and B.1). Lower RD and higher AD values are present on white matter regions (panels A.2-A.3 and B.2-B.3). Diffusion fractional anisotropies show higher values in white matter, particularly for regions corresponding to aligned white matter fiber bundles (panels A.4 and B.4). MD, AD and RD estimates from the DTI model show lower values in comparison to the measures extracted from DKI.

**Figure 3.**
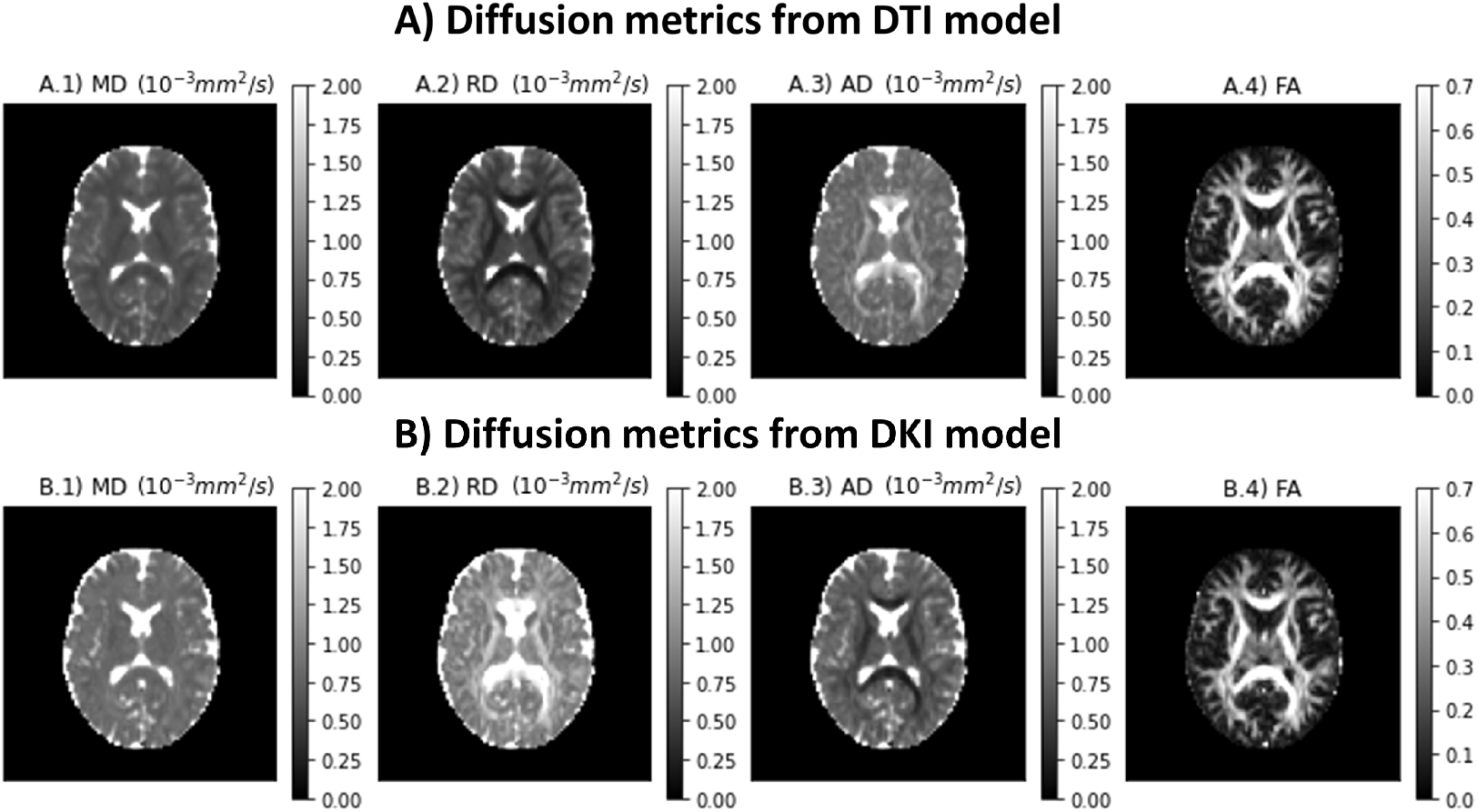
Standard diffusion metrics for a representative axial slice of the CFIN data and extracted from the DTI (A) and DKI (B) models. Left to right panels show the maps of mean diffusivity (A.1 and B.1), radial diffusivity (A.2 and B.2), axial diffusivity (A.3 and B.3), and Fractional Anisotropy (A.4 and B.4)

Different kurtosis-based metrics are shown in Figure 4 for a representative axial slice of the CFIN dataset. MK presents higher intensities in white matter (Fig. 4, panel A). On the other hand, RK shows values higher than the AK (Fig. 4, panels B and C). Although diffusion-weighted data was pre-processed using the PCA denosing and Gibbs unringing algorithms, MK and RK maps present implausible low kurtosis values in deep white matter (e.g. voxels pointed by red arrows in Fig. 4). Mean kurtosis tensor (MKT) estimates show similar contrast to the MK map; however, MKT white matter estimates seem to be less affected by implausible low kurtosis values (Fig. 4, panel D). The kurtosis fractional anisotropy (KFA) map shows smoother profiles (Fig. 4, panel E) when compared to the standard diffusion FA map (Fig. 3, panel D.1), with the exception of sporadic voxels containing high KFA and which corresponds to voxels with implausible negative values in kurtosis estimates (e.g., voxels pointed by the yellow arrows in panels D and E of Fig. 4). Mean signal kurtosis presents similar maps to MK and MKT; however, white matter estimates do not present implausible negative values (Fig. 4, panel F).

**Figure 4.**
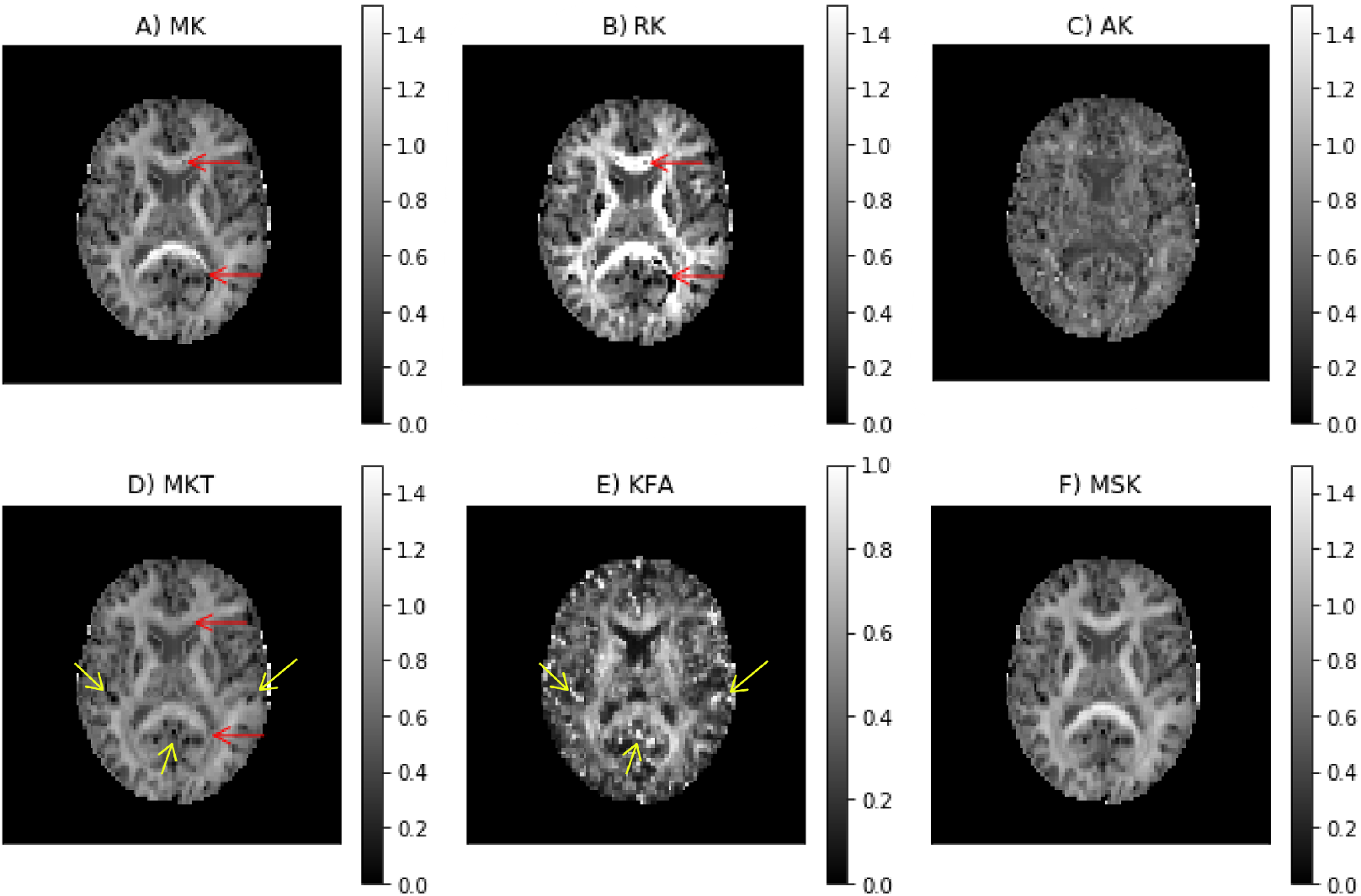
Kurtosis metrics for a representative axial slice of the CFIN data: A) mean kurtosis; B) radial kurtosis C); axial kurtosis; D) mean kurtosis tensor; (E) kurtosis fractional anisotropy; and F) mean signal kurtosis

The results of the two kurtosis-based microstructural models are presented in Figure 5. Axonal water fraction (AWF) and tortuosity estimates from the WMTI model are plotted on well-aligned white matter regions in panels A and B, together with their histograms that reveals similar value ranges to those reported on the original WMTI paper (Fieremans et al., 2011). Panels C and D of Figure 5 show the AWF and intrinsic diffusivity estimates obtained by converting the MSDKI parameters to the SMT2 (MSDKI-SMT2) model parameters (Eq. 22 and Eq. 21). AWF map from the MSDKI-SMT2 model presents a similar contrast than the MSK map (Fig. 4, panel H).

**Figure 5.**
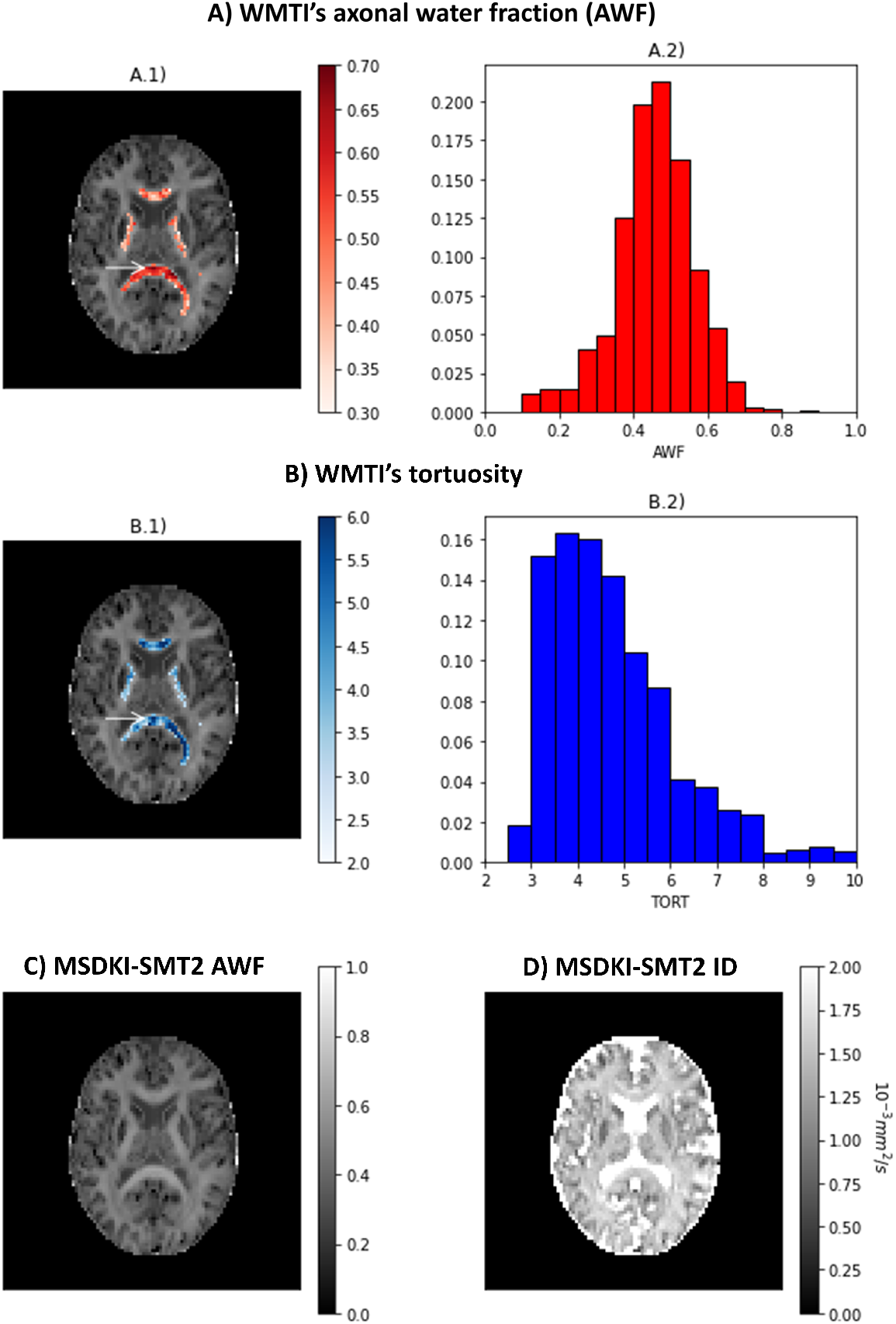
Metrics from the kurtosis-based microstructural models: A) Axonal water fraction (AWF) estimates from the white matter tract integrity model - (A.1) shows the AWF estimates of well-aligned fiber regions overlaid on a top of the mean signal kurtosis image, while (A.2) shows the histograms of AWF estimates for well-aligned fiber regions. B) Tortuosity (TORT) estimates from the white matter tract integrity model - (B.1) shows the TORT estimates of well-aligned fiber regions overlaid on a top of the mean signal kurtosis image, while (B.2) shows the histograms of TORT estimates for well-aligned fiber regions. C) Axonal water fraction (AWF) estimates from the spherical mean technique converted from the MSDKI model. D) Intrinsic diffusivity (ID) estimates from the two-compartmental spherical mean technique converted from the MSDKI model.

### 3.3 Evaluating and comparing the accuracy and reliability of DTI and DKI

We compared the performance of the DKI and DTI model in a large sample: the 789 individuals from the HCP 900-subject release that had completed the entire dMRI measurement. In a k-fold cross-validation analysis, each of the models was fit to 80% of the directions in the data and then the model was used to predict the left-out 20%. This was repeated five times (*k* = 5), such that each measurement was predicted by a model that was fit with this measurement left out. As a measure of model accuracy, we computed the coefficient of determination *R*^2^ between these predictions and the measured data. We found that both DKI and DTI fit the data very accurately: *R*^2^ exceeded 80 for both models in all but one individual (subject 105014), who was an outlier with very low *R*^2^ in both models. In addition, *R*^2^ was consistently higher for DKI than for DTI (this is also true for the outlier subject 105014, not shown in Figure 6), and this difference was statistically significant over the entire sample. This was true both when the DTI model was fit to all of the b-values in the data (Kruskal-Wallis *H* = 1173.4, *p* < 10^−200^), but also when DTI was fit only to the high SNR b-value tier of *b* ≈ 1000*s/mm*^2^, albeit, with a much smaller effect (Kruskal-Wallis *H* = 5.7, *p* < 0.05).

**Figure 6.**
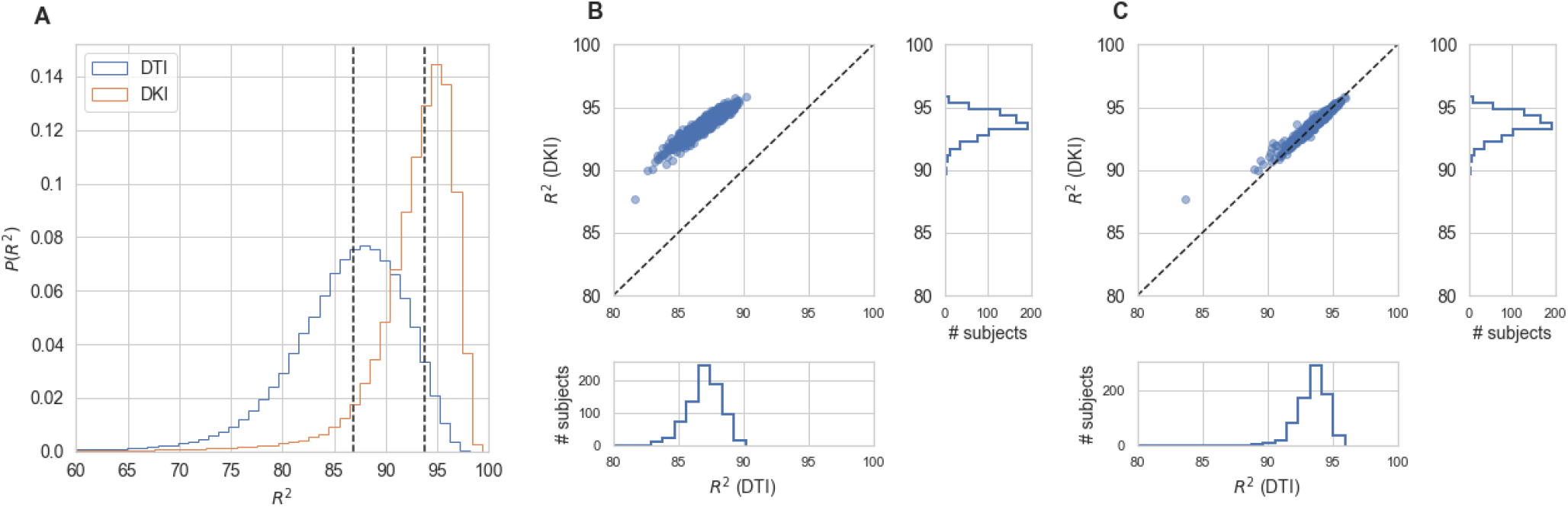
Comparing DTI And DKI accuracy in the Human Connectome Project dataset. Within a typical single individual **(A)**, the distribution of cross-validated *R*^2^ is displayed. Dashed line indicates the median of each distribution. This is also consistent across all of the subjects in the dataset that have measurements of DWI **(B)**. The median cross-validated *R*^2^ in the white matter is always higher for DKI than for DTI. **(C)** This holds, though the effect is substantially smaller, when the DTI model is fit only to data with *b* = 1000*s/mm*^2^

**Figure 7.**
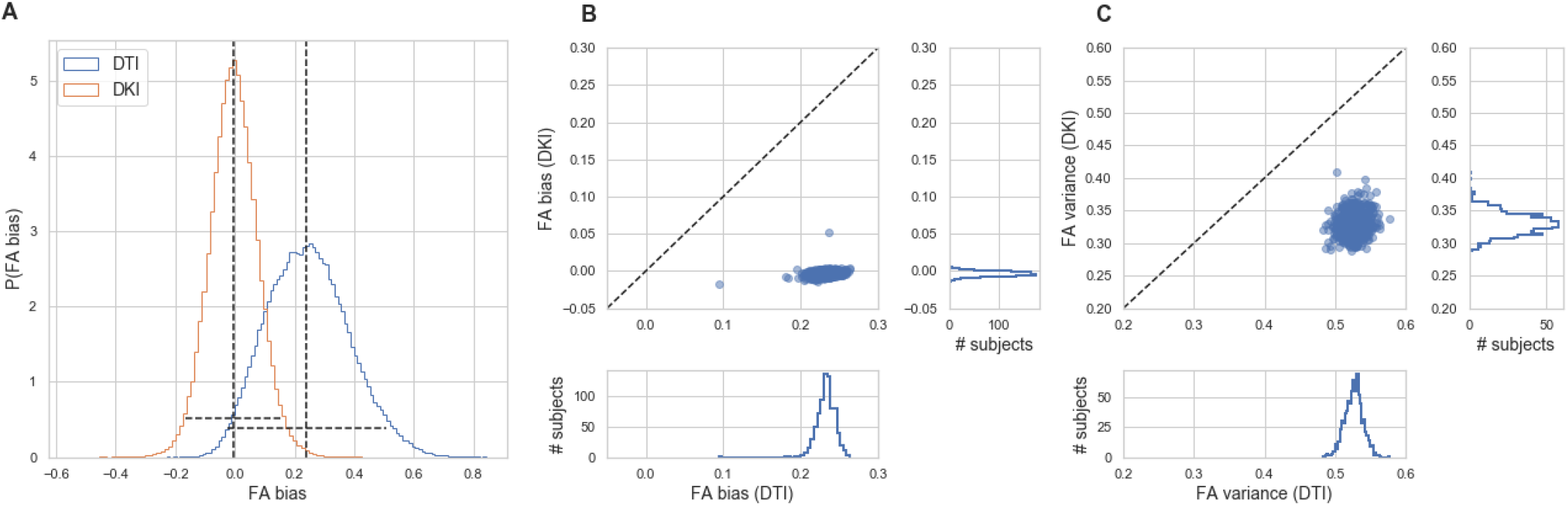
Comparing DTI And DKI variability in the Human Connectome Project dataset. We compared FA for DTI and DKI in different subsets of the data. In each voxel in the white matter, the bias was defined as the modal difference between values of FA between different subsets and the variance was defined as the spread of these values across voxels. **A**For example, within a typical individual, the bias of DKI FA was close to 0, while there was substantial bias in the DTI FA. **B**. The distribution of bias values across subjects is also close to 0 in DKI and mostly non-zero in DTI. **C** Variance is systematically larger in DTI FA than in DKIFA

To evaluate the precision of the two models, we compared the stability in their estimates of the derived FA value and MD. As explained in section 2.1.4, this measure can be computed with both of the models, and their interpretation would be similar, even when values derived from the different models may differ. We used a strategy previously used in comparing these indices derived from the two models in data collected in rat brain (Veraart et al., 2011): data is sub-sampled to different b-values. For DTI, we assessed FA in *b* ≈ 1000*s/mm*^2^ and in a combination of *b* ≈ 1000 + 2000*s/mm*^2^. In DKI, we assessed FA in combinations of *b* ≈ 1000 + 2000*s/mm*^2^ and *b* ≈ 1000 + 3000*s/mm*^2^. We find that FA reliability is consistently higher for DKI than for DTI. This is true both in terms of the bias, and in terms of the variance, both of which are smaller for DKI in all of the subjects in the sample.

## 4 DISCUSSION

Diffusional kurtosis Imaging (DKI) is a straightforward expansion of standard diffusion tensor imaging (DTI). While DTI only quantifies diffusion anisotropy via the diffusion tensor, DKI also quantifies the non-Gaussian properties of diffusion via the diffusional-kurtosis tensor. Although measures derived from DKI do not have a direct mechanistic link to specific biological properties, they are known to be sensitive to differences in microstructural tissue properties. For instance: DKI can provide a better characterization of healthy brain aging alterations in both brain white and grey matter regions than DTI measures (Falangola et al., 2008; Henriques, 2018; Price et al., 2017); it may be used as marker of grey matter neurofilament density in healthy brains (Zhu et al., 2021); it has great potential in discriminating different tumour grades (Lin et al., 2018); it more sensitive to ischemic lesion than DTI measures (Rudrapatna et al., 2014; Hui et al., 2012); it seems to be sensitive to different microstrucutural mechanisms on traumatic brain injury (Zhuo et al., 2012; Grossman et al., 2012; Steven et al., 2014); and it can provide useful markers for tissue degeneration in Parkinson’s and Alzheimer’s diseases (Wang et al., 2011; Struyfs et al., 2015).

Moreover, DKI provides a useful first step in subsequent microstructural model fitting. One well-established DKI-based microstructural model is the white matter tract integrity model (WMTI) Fieremans et al. (2011). While recent studies have raised some doubts about the specificity of WMTI, because of violations of its underlying assumptions (Henriques et al., 2019; Dhital et al., 2018), the metrics derived from WMTI may still be used as sensitive biomarkers, above and beyond the standard DKI measures. For example, previous studies have shown that these measures could potentially be used to reveal some mechanisms behind ischemic stroke lesions (Hui et al., 2012), discriminate different progression stages of Alzheimer’s disease and mild cognitive impairment (Fieremans et al., 2013), and that they shed additional light on brain-behavior correlations in reading abilities(Huber et al., 2019). More recently, the use of the mean signal DKI (MSDKI) model was suggested to overcome confounds of standard DKI and of WMTI due to fiber orientation dispersion Henriques (2018). As shown in this work, MSDKI can be combined with microstructural models as the spherical mean technique models that are general to any direction configuration of microstructural compartments. Taken together, the body of findings on the sensitivity and reliability of DKI, and the range of derived microstructural models provide powerful tools to study human white matter.

Here, we provide a reference implementation of DKI model fitting and related techniques in the DIPY project (Garyfallidis et al., 2014). The implementation is feature complete: it includes several different methods for fitting the basic DKI model and compute derived quantities, as well as relevant extensions to the model: the mean signal DKI (MSDKI) model (Henriques, 2018), WMTI (Fieremans et al., 2011), and the MSDKI-SMT models (Henriques et al., 2019). A reference implementation of these methods that is comprehensive, thoroughly-tested and well-documented is an important accelerant for subsequent scientific research. It provides a proving ground for new methods and a basis for comparison between methods. The fact that DIPY is managed as an open software library, where issues can be publicly reported, discussed and addressed, means that errors can be surfaced by any user of the software. The open-source code means that these issues can be demonstrated and fixed directly by reference to the code itself. This is important for correctness of the implementation, and it also promotes the reproducibility of results obtained using this implementation (Rokem et al., 2018). The modularity and object-oriented design of the software means that statistical procedures that are implemented in one model can be readily translated to other models. For example, we demonstrate here the use of cross-validation for model evaluation (Rokem et al., 2015). The DIPY application programming interface provides uniform methods for resampling such that the model is fit in some directions and predicted in other directions. This software architecture is extensible. We have already used this fact to extend DKI. But it also means that others can rely on the architecture to build future developments.

### 4.1 Findings

To demonstrate the utility of our software, we analyzed several different datasets. Numerical simulations were first used to demonstrate the sensitivity of the DKI method and the software in known microstructural configurations. Particularly, based on these simulations, we demonstrate that DKI does not only provides a quantification of non-Gaussian diffusion but also decouples non-Gaussian diffusion effects from standard diffusion tensor metrics - a reason why DKI diffusion tensor estimates more closely match their ground truth estimates than the DTI tensor estimates 1. Simulations were also used to illustrate that, while systems comprising different components with distinct diffusivities and configurations can present very similar diffusivities, kurtosis estimates can help distinguish them by providing information on diffusion heterogeneity 2. Our simulations also reproduce the kurtosis geometries exploration of (Henriques et al., 2015) which revealed that maximum kurtosis values are present perpendicular individual fibers even in crossing configurations, and confirmed that MSK estimates are invariant to the directional configuration of tissue compartments.

We used the CFIN dataset to provide examples of the contrasts provided by DKI. This is a dataset that is directly and openly accessible to anyone through the DIPY dataset interface, so the figures using this data can be reproduced with code that we provide (and is also provided as part of the DIPY documentation, both for standard DKI^2^ and for MSDKI ^3^). This data is used in tandem with supporting methods that address some of the limitations of the method (see below): to produce the maps shown in Figures 3 and 4, we used both denoising (Veraart et al., 2016b) and Gibbs ringing removal Kellner et al. (2016); Henriques (2018), both implemented in DIPY. Based on this sample dataset, we demonstrate the typical contrasts of Mean, axial and radial diffusivities and kurtosis. We illustrate that MK, MKT and MSK present similar contrasts (consistent to what was reported by (Hansen et al., 2013)), but MSK was shown to be more robust to image noise and artifacts. Moreover, we confirmed that KFA provides different contrast information than standard FA measures (Glenn et al., 2015b; Hansen and Jespersen, 2016b; Hansen, 2019).

The CFIN dataset was also used to illustrate the estimates obtained from the implemented kurtosis-based microstructural models 5. Axonal volume fraction and extracellular tortuosity estimates from the white matter tract integrity model (WMTI) showed similar value ranges than reported on the original WMTI paper (Fieremans et al., 2011). As one may expect from the theory (e.g., equation 22), we also highlighted that the axonal water fraction maps obtained for the MSDKI-SMT2 model provide similar contrast to MSK. Since previous studies showed that SMT2 assumptions do not properly represent biological tissues (Henriques et al., 2019), AWF should not be interpreted as accurate biophysical estimates of axonal water fraction. Instead, it could be a useful normalized version of MSK scaled in a range between 0 and 1.

Finally, we analyzed a large, openly available dataset provided by the Human Connectome Project (Sotiropoulos et al., 2013; Glasser et al., 2016). In this dataset, we demonstrated the comparison of the DKI model and DTI model in cross-validation (Rokem et al., 2015). We found that DKI consistently fit the data more accurately than DTI. In addition, the FA derived from DKI showed more stability across different sub-samples of the data. Considering these two facts, we conclude that in this dataset, FA and other metrics should be computed using the DKI model. As made clear above, in addition to its higher accuracy and precision, the DKI model provides a wealth of additional information. These findings are important in the context of the HCP dataset, as this data is likely to be analyzed by many other researchers. In addition, several efforts are currently underway to collect similar large-scale datasets with multiple diffusion weighting values (e.g., (Jernigan et al., 2018) and (Alexander et al., 2017)), and similar conclusions may apply in these datasets as well.

### 4.2 Related work

There are several other software implementations of DKI that are available. These include implementations in the DKE software Tabesh et al. (2011), available through NITRC ^4^ and ExploreDTI (Leemans et al., 2009). But neither of these software projects is open-source or provided through an OSI-approved license, limiting their broad use. In addition, they both require the proprietary Matlab software platform. Another Matlab-based software that provides open-source and OSI-approved licensed software is DESIGNER (Ades-Aron et al., 2018b). Finally, the recent version of the mrtrix software (Tournier et al., 2019) also includes an implementation of DKI estimation and calculation of DKI-based metrics. The software presented here adds to these software projects in that it includes Python-based implementations that are, on the one hand, completely open-source and gratis for use in any context. On the other hand, using the Python programming language enhances the readability of the code and its understanding by researchers from many different backgrounds (e.g., relative to scientific software implemented in compiled languages, such as C/C++). The software is specifically designed to enable extensions and further developments on top of the existing methods. These design considerations have allowed our development team to expand the original implementation of DKI to include many different additional methods, described above.

### 4.3 Limitations and Future work

As we highlight in this work, DKI can provide more accurate diffusion estimates than DTI in addition to measures of diffusional kurtosis. However, it is important to note, fitting DKI requires data from multi-shell b-values, and thus it cannot be used to analyze data acquired using single non-zero b-value acquisitions. Moreover, since it involves the estimation of a larger number of parameters than DTI, DKI can provide less precise (i.e. noisier) estimates. For instance, as illustrated in Fig 4, standard DKI maps can typically provide implausible negative estimates “black voxels”. These negative kurtosis estimates are not associated with fitting implementation errors but with effects of thermal noise and may predominantly appear in regions where diffusivities are low (Tabesh et al., 2011; Henriques, 2012; Kuder et al., 2012; Veraart et al., 2013). Kurtosis in these regions present very low robustness, since it is very hard to capture non-Gaussian information due to the low decays of that region in radial direction. This is the case of well-aligned fibers that have low diffusivities in radial direction. Therefore, in white matter, implausible kurtosis typically appears in regions such as in the corpus callosum midsagittal plane on both RK and MK mpas. Implausible negative kurtosis estimates can also originate from effects of different image artefact such as Gibbs Ringing artefacts as explained by Veraart et al. (2016a) and Perrone et al. (2015). To suppress effects of noise and artefact, pre-processing techniques such as the Marchenko–Pastur PCA denoising and Fourier sub-voxel shift Gibbs artefact correction algorithm (Veraart et al., 2016b; Kellner et al., 2016) are used. Other pre-processing techniques to suppress artifacts available in DIPY, include the threshold-based PCA denoising (Manjón et al., 2013), the non-local means denoising filter (Coupé et al., 2011) and the self-supervised denoiser Patch2Self (Fadnavis et al., 2020). Particularly, Fadnavis et al. demonstrated that Patch2Self outperforms current implementations of low-rank method approximations, such as MP-PCA (Fadnavis et al., 2020). (which has denoising performance similar to the PCA denoising algorithms, but with less processing time demands (Fadnavis et al., 2020)). Implausible kurtosis estimates can also be mitigated by excluding data outliers (Chang et al., 2005; Tax et al., 2015)), imposing constraints (Tabesh et al., 2011; Kuder et al., 2012), and readjusting b-value=0 data points (Zhang et al., 2019). Alternatively, if only interested in isotropic kurtosis measures, one could opt to use of the MSK estimates which provide more precise quantification of non-Gaussian diffusion in low-diffusivity regions (c.f. Fig. 4). In addition, MSK can be used to regularize the full DKI providing more robust kurtosis tensor derived metrics (Henriques et al., 2021)). In future studies, DIPY can provide a useful framework for the comparison of all these different noise and artefact suppression techniques and for the development of novel strategies.

As with other techniques based on the diffusion-weighted signal cumulant expansion, DKI estimates may be biased by high-order-term effects not considered by the expansion truncation (Chuhutin et al., 2017). While higher-order-effect in diffusion tensor metrics are minimized by DKI (c.f. Fig. 1, panel B), these can introduce the deviations between kurtosis tensor estimates and their ground truth values. Despite these accuracy issues, fitted kurtosis tensors still present a fair description of the 3D information of the kurtosis tensor as shown in panel C of Fig. 1. Although DKI may provide robust information from microstructural model fitting (Fieremans et al., 2011; Jespersen, 2018; Jespersen et al., 2018), it is important to note that the higher-order-term biases on DKI can propagate to DKI-based microstructural estimates. Therefore, in future studies it will be of interest to combine our implemented strategies to the full analytical derivations of different microstructural models. For instance, the more robust DKI-based estimates could be used as the initial guess estimates for the more complex non-linear fitting procedures required by some current microstructural models. Moreover, we expect that our DKI microstructural models could be expanded to remove some of the model constraints assumed by WMTI and SMT2 models (e.g. incorporating tissue dispersion and removing the lower intra-cellular diffusivity constraint of the WMTI model (Jelescu et al., 2016; Novikov et al., 2018b; Jespersen, 2018; Jespersen et al., 2018).

While DKI can provide additional information to DTI, the non-Gaussian diffusion information provided by single diffusion encoding (SDE) multi-shell acquisitions can originate from multiple sources, and thus limiting kurtosis specificity (Szczepankiewicz et al., 2016; Henriques et al., 2020a). For example, kurtosis decreases in gradual degeneration processes (e.g., healthy ageing, Alzheimer’s and Parkison’s diseases, multiple sclerosis) are typically attributed to a reduction in individual fibers’ diffusion anisotropy (Falangola et al., 2008; Wang et al., 2011; Struyfs et al., 2015; Andersen et al., 2020). On the other hand, kurtosis increases in more abrupt damaged microstructural tissues (e.g., ischemia, traumatic brain injury) can occur due to cytogenic and vasogenic effects (Hui et al., 2012; Zhuo et al., 2012). To increase the specificity of kurtosis, several studies have used advanced diffusion encoding sequences to decouple different kurtosis sources (e.g. (Westin et al., 2016; Topgaard, 2017; Szczepankiewicz et al., 2020, 2016; Henriques et al., 2020a,b)). Due to the high potential of these techniques, procedures for reconstructing diffusion-weighted data acquired from advanced diffusion sequences are currently being incorporated in DIPY.

The present work focused on the use kurtosis to probe microstructural and biophysical properties of the tissue in different locations within the white matter. However, DKI can also provide information to identify the trajectories of white matter bundles through the brain, connecting local or remote parts of the brain to each other through computational tractography (Lazar et al., 2008; Jensen et al., 2014; Henriques et al., 2015; Glenn et al., 2015a, 2016). To implement the DKI-based tractography in DIPY we would require an implementation of an analytical solution that relates the diffusion and kurtosis tensors to the tissue orientation distribution function (ODF). This will be included in future releases of DIPY. For the time being, DIPY provides many other methods to compute ODFs, including from DTI, as well as from constrained spherical deconvolution (Tournier et al., 2007) (including a multi-shell variant of this method (Jeurissen et al., 2014)).

## AUTHOR CONTRIBUTIONS

RNH led the implementation of the software. MC, MM, SK, EG and AR contributed to, and reviewed the software implementation. JY and EH tested the software in real datasets and contributed to the implementation of microstructural modeling API. JK conducted analysis HCP data. RNH and AR conducted simulations, analyzed data and wrote the manuscript, with feedback from all other authors.

## ACKNOWLEDGMENTS

Data were provided in part by the Human Connectome Project, WU-Minn Consortium (Principal Investigators: David Van Essen and Kamil Ugurbil; 1U54MH091657) funded by the 16 NIH Institutes and Centers that support the NIH Blueprint for Neuroscience Research; and by the McDonnell Center for Systems Neuroscience at Washington University.

## DATA AVAILABILITY STATEMENT

The datasets analyzed for this study are openly available. The CFIN dataset is available through https://datadryad.org/stash/dataset/doi:10.5061/dryad.9bc43 and can also automatically be downloaded using DIPY’s data fetcher module. The Human Connectome Project dataset can be accessed via https://www.humanconnectome.org/.

## Footnotes

1 http://gsoc2015dipydki.blogspot.com/2015/07/rnh-post-8-computing-perpendicular.html

2 https://dipy.org/documentation/latest/examples_built/reconst_dki/

3 https://dipy.org/documentation/latest/examples_built/reconst_msdki

4 https://www.nitrc.org/projects/dke/

